# Visualization of *Bacillus subtilis* Spore Structure and Germination using Quick-Freeze Deep-Etch Electron Microscopy

**DOI:** 10.1101/2024.03.04.579732

**Authors:** Kiran Jalil, Yuhei O. Tahara, Makoto Miyata

## Abstract

Bacterial spores, known for their complex and resilient structures, have been the focus of visualization using various methodologies. In this study, we applied quick-freeze and replica electron microscopy techniques, allowing observation of *Bacillus subtilis* spores in high-contrast and three-dimensional detail. This method facilitated visualization of the spore structure with enhanced resolution and provided new insights into the spores and their germination processes. We identified and described five distinct structures: (i) hair-like structures on the spore surface, (ii) spike formation on the surface of lysozyme-treated spores, (iii) the fractured appearance of the outer spore cortex during germination, (iv) potential connections between small vesicles and the core membrane, and (v) the evolving surface structure of nascent vegetative cells during germination.

## Introduction

The phylum Firmicutes is divided into two categories: aerobic *Bacillaceae* and anaerobic *Clostridia*. Both groups produce structurally similar spores, exemplified by pathogenic *Bacillus anthracis* and *Clostridium difficile*, along with environmental *B. subtilis* [1–3] These spores exhibit remarkable resistance and survivability compared to their vegetative states. This is attributed to their protective outer coat layers, the peptidoglycan (PG) layer of the spore cortex, reduced inner membrane permeability, high concentrations of dipicolinic acid with divalent cations, and specialized low-molecular-weight protective proteins. Such features are crucial for maintaining low core water content and safeguarding spore DNA [4–6].

To date, it is known that the *B. subtilis* spore comprises six layers (Fig. 1). The outermost crust consists of glycoproteins, which likely contribute to the spores’ hydrophobicity, adhesion, dispersal capabilities, and resistance to environmental stresses [7–9]. The second layer (the rodlet layer) has a distinct surface pattern [7, 10, 11]. The cortex exhibits a unique PG structure characterized by reduced peptide cross-linking, achieved through the removal of peptide side chains and the conversion of muramic acid to muramic lactam. A few sections resemble vegetative cell walls with glycan chains. The inner membrane that encloses the core plays an essential role in resistance and germination [6, 12]. The core itself contains a balanced mixture of <u>Ca</u>2+ and <u>d</u>i<u>p</u>icolinic <u>a</u>cid (CaDPA), ribosomal proteins, and certain lytic enzymes [13, 14].

**Figure 1.**
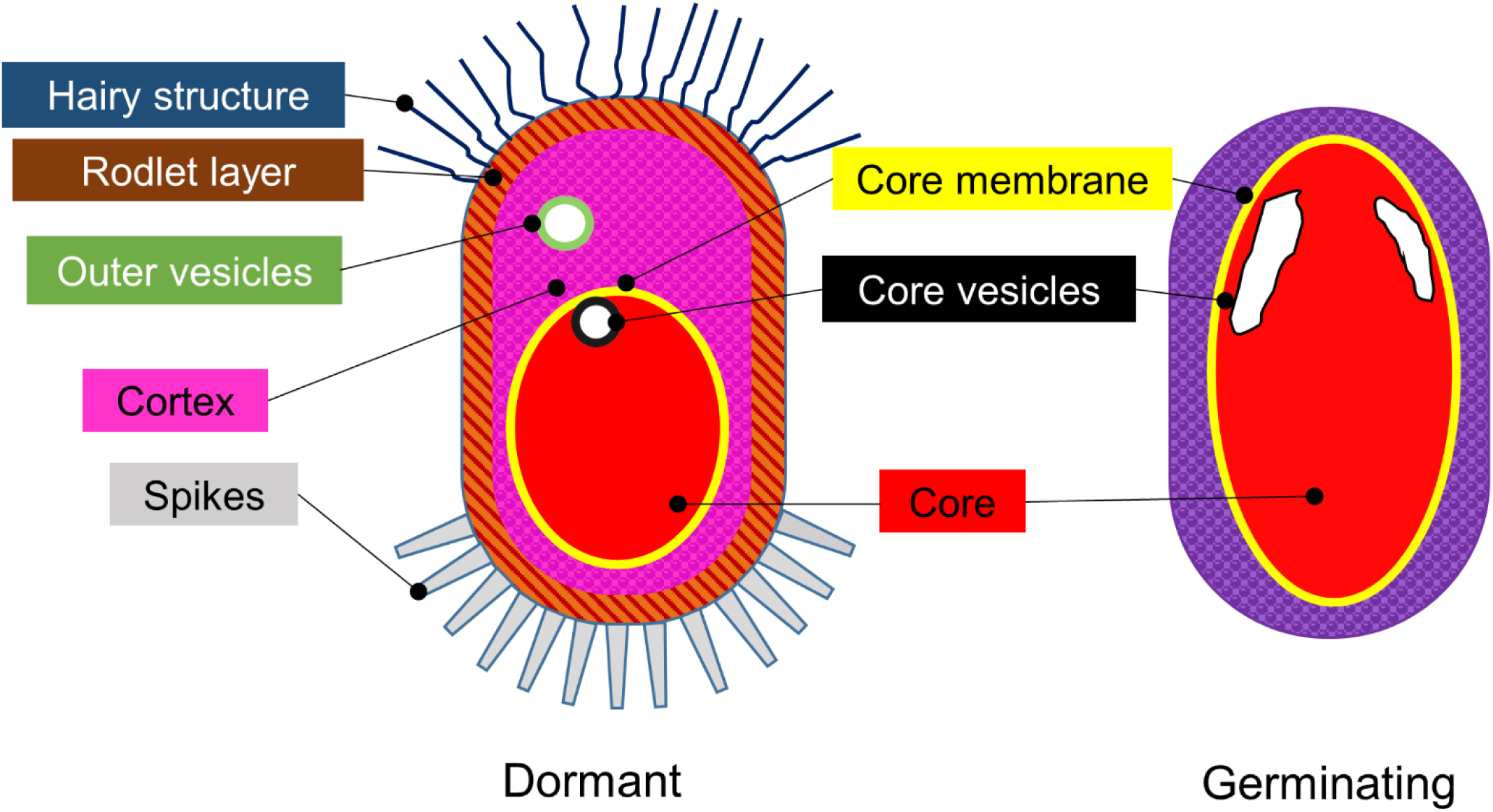
Schematic illustrations of spore structures. Left: Dormant spore. Right: germinating spore. Hair-like structures are observed on the dormant spore.

Upon exposure to germinants such as amino acids and peptides, receptors on the inner membrane are activated. This leads to the release of certain enzymes and CaDPA from the core, resulting in the hydration of the core. The inner membrane and core remain intact, eventually transforming into a vegetative cell [13, 14].

In this study, we used the quick-freeze deep-etch electron microscopy (QFDE-EM)_technique to visualize the entire structure of *B. subtilis* spores and their germination processes at high resolution. Introduced in 1979 to observe the vesicle release of neurotransmitters in nerve impulses in frogs, this technique differs from traditional “freeze and replica electron microscopy (EM)” by avoiding chemical fixation and utilizing surface exposure through deep etching [15, 16]. The specimens were rapidly frozen on a metal block cooled with liquid helium or nitrogen, achieving fixation much faster than using chemical methods. Subsequently, the specimens were fractured and etched, followed by platinum shadowing. This shadowing enhances image contrast, enabling the capture of detailed images without the need for image averaging. Moreover, it is particularly suited for visualizing surfaces with low density and flexible structures and the internal structure of specimens at high resolution, making it invaluable for microbiological studies focused on surface structures [15–17].

## Material and methods

### Strain, media, and culture conditions

*B. subtilis* 168 CA was cultured in Luria-Bertani (LB) medium. To induce spore formation, the cultured cells were transferred to modified Schaeffer’s medium [18]. Spores were harvested after 12 h at 4°C, followed by two rounds of washing with ice-cold water to eliminate vegetative cells [19].

### Lysozyme treatment

Spores (50 mg dry weight) were suspended in 1 mL of phosphate-buffered saline (PBS; 75 mM sodium phosphate, pH 7.3, 68 mM NaCl) containing 1 mg of lysozyme [20] and incubated at 37°C for 2 h with shaking. Post-incubation, spores were pelleted via centrifugation at 8,000 × g for 3 min at 25°C, then suspended in 1 mL of a solution containing 0.05% SDS, 10 mg/mL RNase, and 10 mg/mL DNase, and incubated for 1 min at 25°C. Following two washes with pure water, the spores were suspended in 0.5 mL of pure water.

### Strain, media, and culture conditions for vegetative cells

Vegetative *B. subtilis* cells were cultured in an LB medium. Next, 100 µL of this culture was inoculated into fresh LB medium and incubated with shaking for 2 h at 37°C.

### Isolation of vegetative PG

Five mL of the vegetative cell culture was centrifuged at 10,000 × g for 5 min at 25°C. The cell pellet was suspended in 1 mL of PBS, transferred to a 1.5 mL tube, and centrifuged again using the same conditions. The pellet was then treated with 0.5 mL of 10% SDS and incubated at 96°C for 3 h, followed by incubation at 25°C for 20 min and centrifugation at 20,000 × g for 30 min at 25°C. After discarding the supernatant, the pellet was treated with 10 µL of 10 mg/mL chymotrypsin at 37°C for 2 h, washed, and finally resuspended in 50 µL of pure water [21]

### Isolation of spore PG

Spores underwent a modified PG isolation procedure [22], including boiling in 50 mM HEPES buffer (pH 7.8) containing 4% SDS, 30 mM DTT, and 2 mM EDTA for 12 h, followed by incubation at 37°C for 45 min. After multiple washes, the spores were treated with 2 mg proteinase K at 70°C for 1 h, washed, and then treated with lysozyme at 37°C for 12 h.

### Germination

Germination of spores was triggered by heating at 70°C for 30 min and introducing a germination medium containing 10 mM L-alanine, 1 mM D-glucose, 1 mM sodium chloride, and 25 mM HEPES buffer (pH 7.8), according to procedures described in a previous study [23] with slight modifications.

### Microscopy

The spores were prepared for microscopy using multiple washes and were resuspended in pure water. The concentration was adjusted to 20 times the original culture density for observation by phase-contrast optical microscopy and negative staining EM. Quick-freeze deep-etch EM was performed as previously described in earlier studies [17, 21, 24].

## Results

### Surface structures

In this study, QFDE-EM was used to examine the surface structures of isolated spores. These spores were combined with mica flakes, rapidly frozen, fractured, and etched. Their replicas were subsequently analyzed (Fig. 2). Observations revealed that the density of spores adhering to the mica flakes was equivalent to that observed using optical microscopy (Fig. 2a, b). Furthermore, the measurements of the spore dimensions using QFDE-EM agreed with those obtained using optical microscopy. A comprehensive layer of hair-like structures enveloping the entire spore surface was identified (Fig. 2c), with diameters ranging between 8 and 14 nm (Fig. 2d). Considering the application of a 2 nm-thick platinum coating using QFDE-EM, the actual diameters of these structures were deduced to be between 4 and 10 nm [19]. These hairlike structures varied in length and extended outward from the edges of the spores.

**Figure 2.**
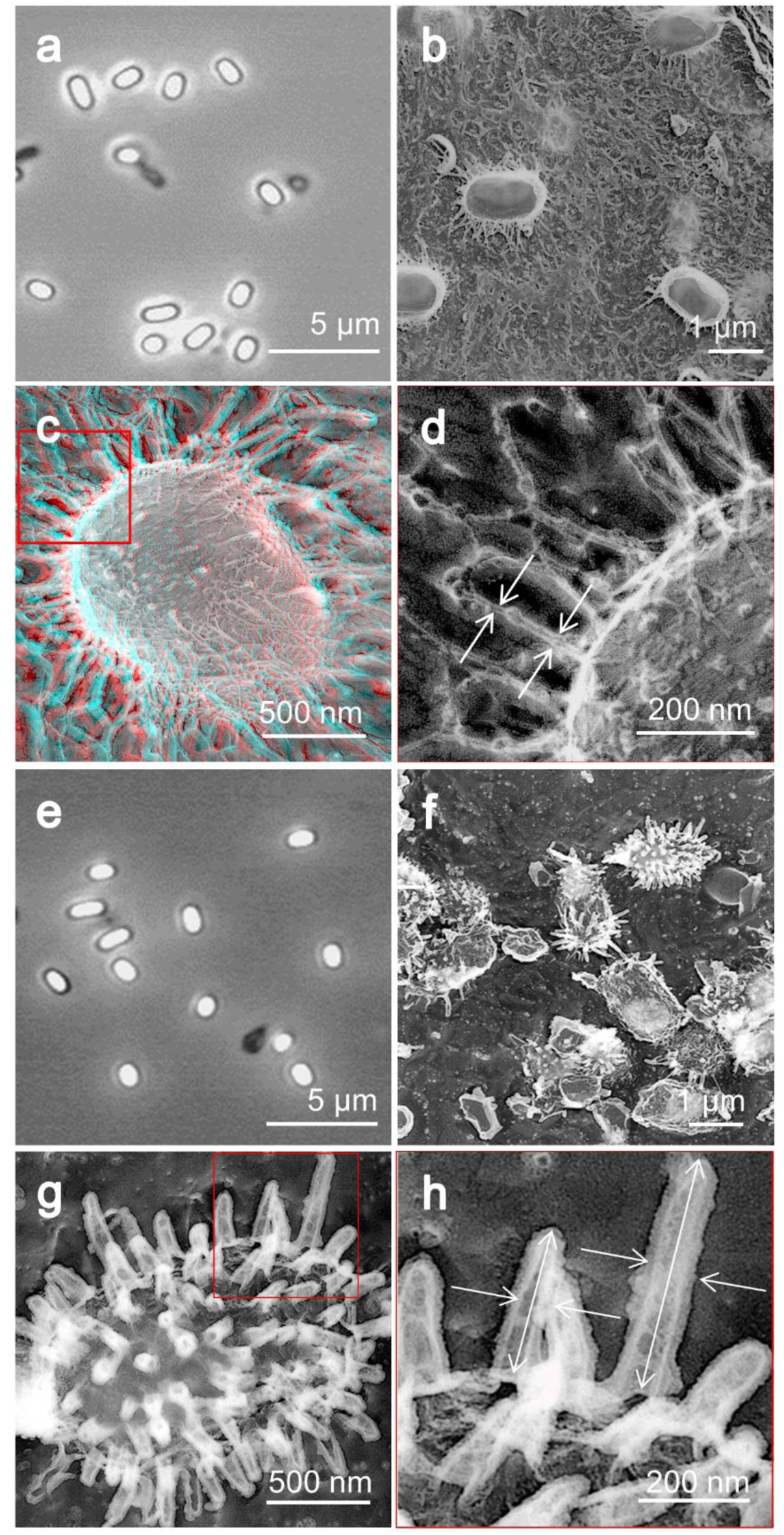
Surface structures. (a) Phase-contrast microscopy of untreated spores. (b, c, d) Quick-freeze deep-etch (QFDE) EM. (c) is a stereoscopic view observed using a pair of glasses with red filter on the left side and blue on the right side. (d) Magnified image of the boxed area in (c). The diameter of the hair-like filament was measured as shown by white arrows. (e) Phase-contrast microscopy of spores treated by lysozyme. (f, g, h) QFDE-EM of lysozyme-treated spores. (h) Magnified image of the boxed area in (f). The lengths and anaglyph diameters of the spikes were measured as shown by the white arrows.

To isolate spores more effectively, we treated them with 1 mg/mL of lysozyme at 37°C for 2 h with agitation. This treatment did not alter the appearance of the spores under a phase-contrast microscope (Fig. 2e). However, the QFDE-EM analysis revealed a transition from hair-like to spike-like surface structures (Fig. 2f). These spike-like formations, ranging from 160 to 510 nm in length, were uniformly distributed across the spore surface at a density of 40–50 spikes per square micrometer (Fig. 2g). Based on 35 observations, the spikes were observed to taper towards the distal end, with an average central diameter of 63.3 ± 13.9 nm (Fig. 2h). The identified structures are shown in a schematic diagram (Fig. 1).

### Internal layers

In the analysis of deeply etched replicas, exposure of a rodlet coat layer situated beneath the hair-like structures was occasionally observed (Fig. 3a). This layer is characterized by uniformly aligned rods, with intervals between these rods determined using fast Fourier transform (FFT) analysis to average 7.8 nm ± 0.44 (Fig. 3b, c).

**Figure 3.**
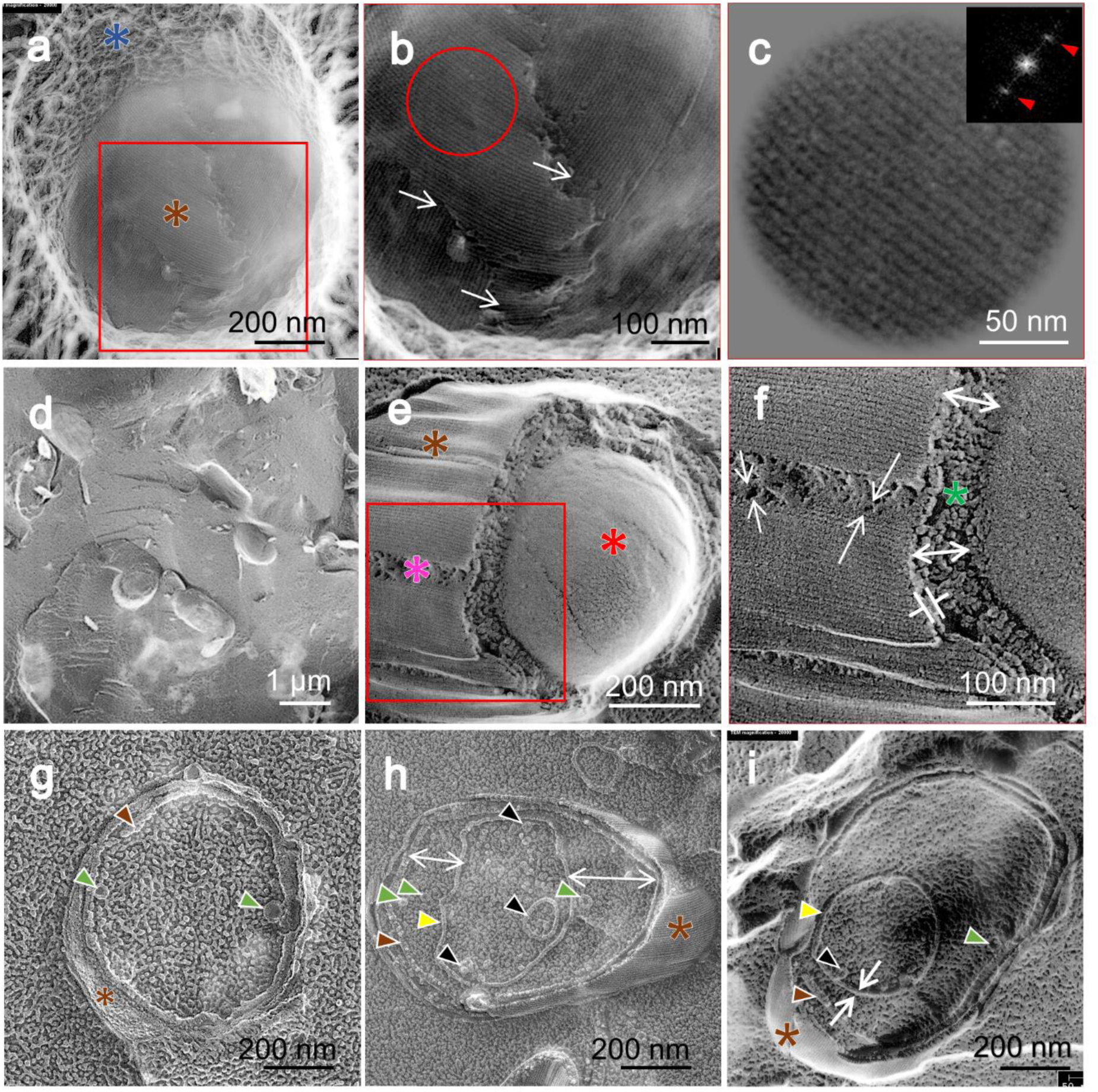
Dormant spores are visualized by quick-freeze deep-etch EM. (a) A slightly fractured spore features a hair-like structure on the surface (blue asterisk) and a rodlet coat layer (brown asterisk). (b) Magnified image of the boxed area in (a). The rodlet coat layers overlap, as marked by white arrows. (c) Magnified and fast Fourier transform (FFT) images of the circle area in (b). The FFT image is characterized by spots marked by red triangles. (d) Field image of fractured spores without etching. (e) Fractured image of a spore. The rodlet layer (brown asterisk), cortex (pink asterisk), and core surface (red asterisk) are shown. (f) Magnified image of the boxed area in (e). White arrows show dimensions. A sectional view of the cortex is marked by a green asterisk. (g, h, i) Sectional image of spore. The rodlet layer (brown asterisk, brown triangles), outer vesicle (green triangles), core membrane (yellow triangles), and core membrane vesicles (black triangles) are shown.

To further investigate the internal structures, a quick-freeze fracture EM without Etching (QFF-EM) technique was employed (Fig. 3d). Spore mixtures containing mica flakes and glycerol at a final concentration of 30% were frozen. The use of glycerol served to minimize eutectic artifacts. This method revealed a segment of the rodlet layer, estimating its thickness to range between 7.6 and 14 nm (Fig. 3e, f). Under this layer, a sparser cortex was discernible; its thickness varied from 49 to 95 nm (Fig. 3e, f), and the surface of the core membrane was visible (Fig. 3e, f).

Occasionally, deep fracturing in the presence of glycerol produced “sectioned” images (Fig. 3g, h), unveiling fragments of another rodlet layer beneath the initial one. This underlying layer had an average thickness of 19.8 nm, and at times, a core membrane with an apparent thickness of 14.3 nm was observed (Fig. 3h, i). The core membrane is not discernible in Fig. 3g, likely because it may not have been exposed during the fracturing process. The sectioned images highlighted membrane vesicles, identified as “sCMMs” (sub-CM membranes) [14], both inside and outside the core, with diameters ranging from 20 to 90 nm (n = 12) and 20 to 60 nm (n = 12), respectively.

### Effect of lysozyme on the isolated cortex

Initially, we isolated the PG layer of vegetative cells via treatment with SDS, followed by enzyme treatment (Fig. 4a–c). Under negative-staining EM, a sack-like structure was observed (Fig. 4b), and a mesh-like structure was discernible using QFDE-EM (Fig. 4c, m). Subsequently, we aimed to isolate the spore PG by treating the isolated spores with 4% SDS and 30 mM DTT at 95°C for 12 h. This treatment resulted in cells appearing darker, as observed under phase-contrast microscopy (Fig. 4d, g), and imparted a slightly rough texture to the cell surface, as observed using QFDE-EM (Fig. 4n).

**Figure 4.**
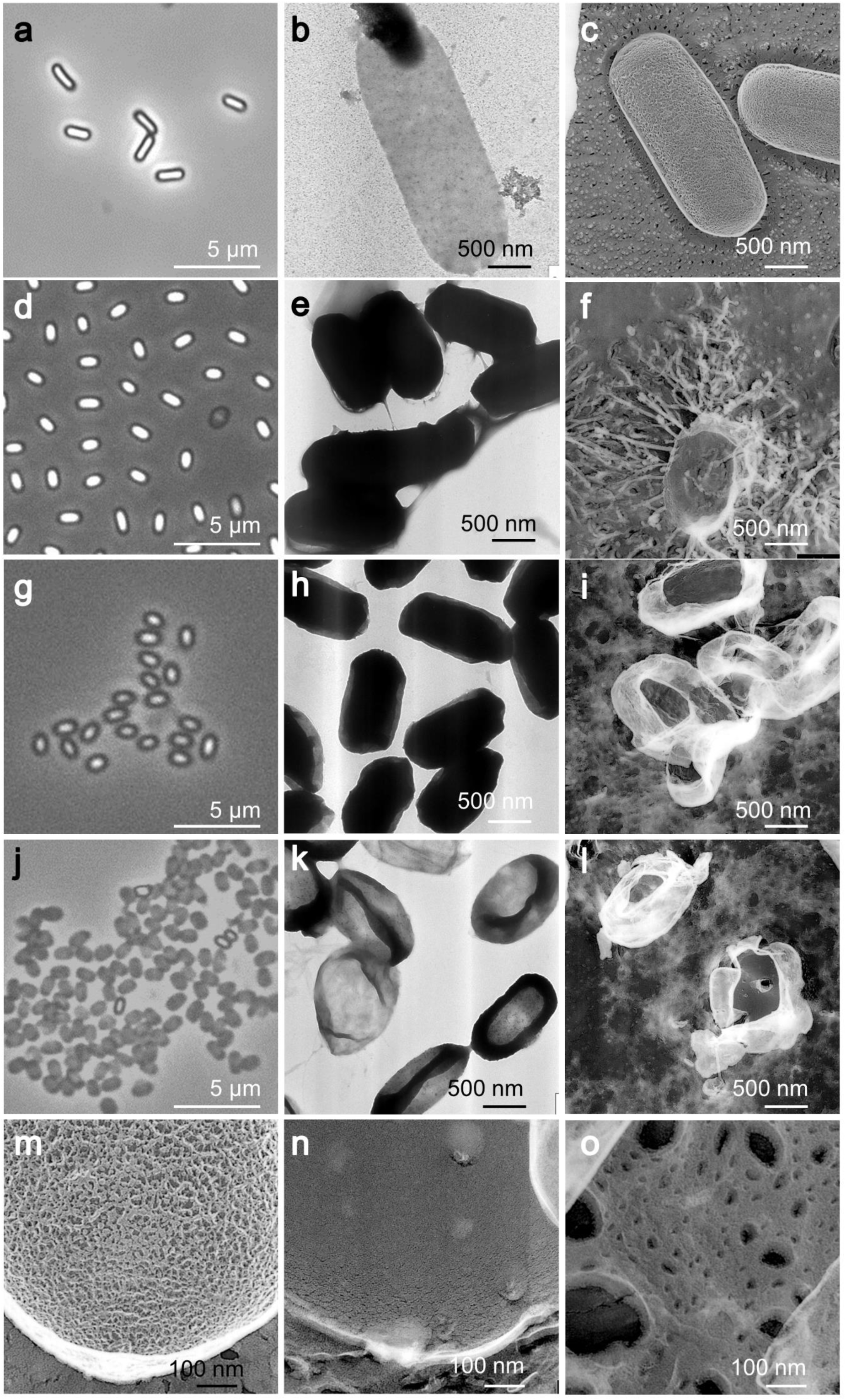
Effect of lysozyme on the isolated cortex. (a) Phase-contrast microscopy of vegetative cells treated with SDS. (b) Negative staining EM of treated vegetative cells. (c) Quick-freeze deep-etch EM of vegetative cells. (d) Phase-contrast microscopy of untreated spores. (e) Negative staining EM of untreated spores. (f) Quick-freeze deep-etch EM of untreated spores. (g) Phase-contrast microscopy of spores treated with SDS, DTT, and proteinase K. (h) Negative staining EM of treated spores. (i) Quick-freeze deep-etch EM image of treated spores. (j) Phase-contrast microscopy of spores treated with SDS, DTT, and proteinase K and then with lysozyme. (k) Negative staining EM of spores additionally treated with lysozyme. (l) Quick-freeze deep-etch EM of spores additionally treated with lysozyme. (m) Magnified image of vegetative cells treated with SDS. (n) Magnified image of spores treated with SDS, DTT, and proteinase K. (o) Magnified image of spores additionally treated by lysozyme.

In contrast to the significant removal of content from the PG layer of vegetative cells, the removal from spores was not readily apparent, as indicated by negative staining EM (Fig. 4h). Then, further treatment of the isolated structures with 0.3 mg/mL lysozyme at 37°C for 12 h was employed to cleave the β (1–4) linkages between N-acetylmuramic acid and N-acetylglucosamine in the PG layers [17, 25]. This enzymatic action resulted in a noticeable transformation visible under phase-contrast microscopy (Fig. 4j). Negative staining EM indicated a significant reduction in cell image density, suggesting that sac structures similar to those of vegetative cells were isolated (Fig. 4k). When subjected to QFDE-EM (Fig. 4l, o), these sac structures displayed pores ranging from 6 to 120 nm in diameter.

### Germination

We then visualized the germination process. The isolated spores were activated by heating at 75°C for 30 min and subsequently incubated in a germination buffer containing 10 mM alanine, 1 mM NaCl, 1 mM glucose, and 25 mM HEPES buffer (pH 7.8) at 37°C. Subsequently, the spores were recovered for structural analysis.

After being incubated for 45 min in the germination buffer, noticeable changes on the cell surface were observed, including cracks in the rodlet layer and exposure of the cortex in certain spores (Fig. 5a–g). However, the hair-like structures remained intact in certain spores (Fig. 5c). To examine the internal structure of the spores at the onset of germination, we subjected the germinating spores to QFF-EM (Fig. 5h–j). In most of the germinating spores, the area delineated by a membrane, identified as the germ cell area, occupied the entire space (Fig. 5h). Additionally, small fragments beneath the germ cells were identified (Fig. 5i, j), possibly originating from vesicles observed in dormant cells, as such vesicles were not observed at this stage.

**Figure 5.**
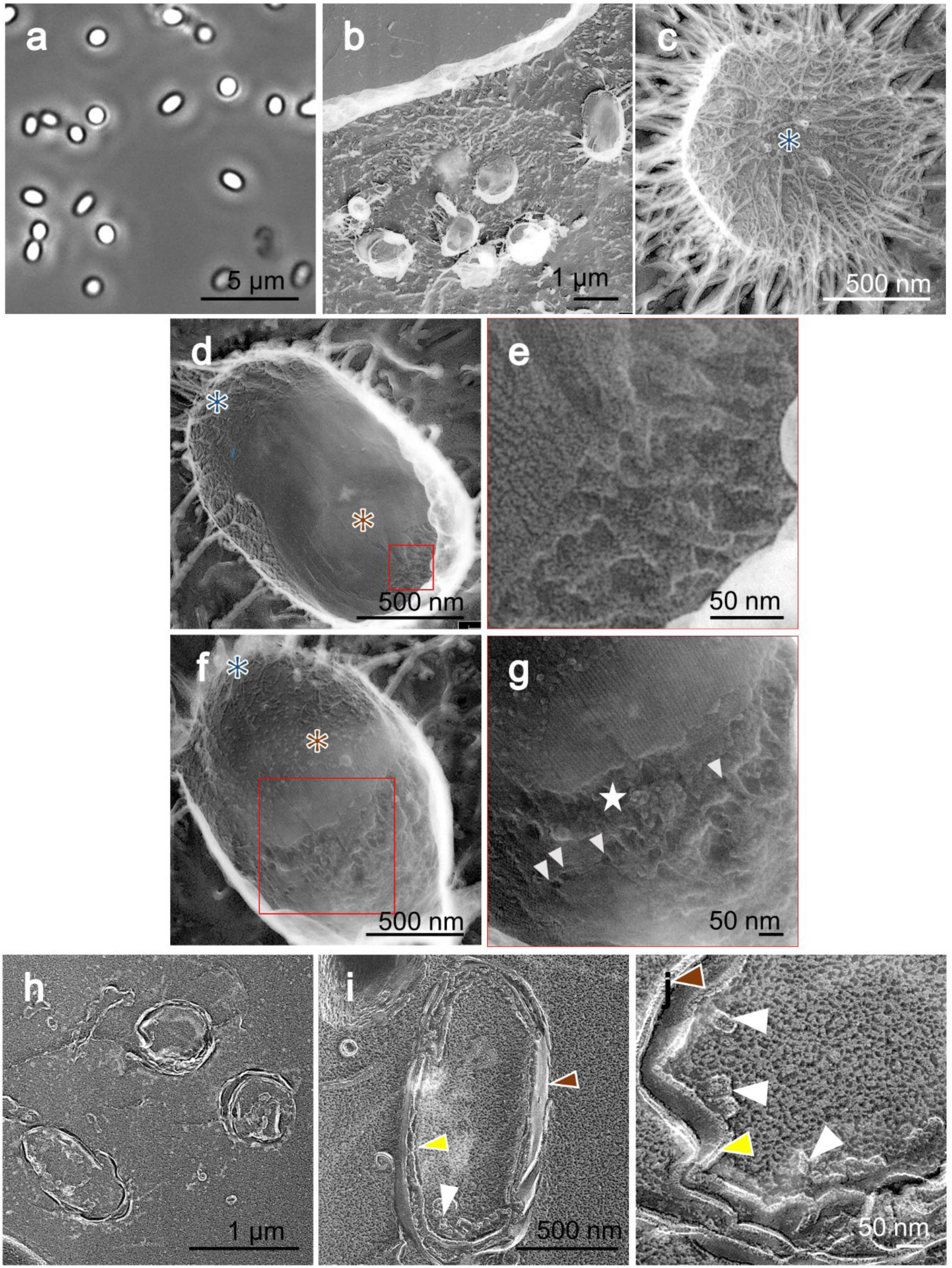
Germinating spores after 45 min incubation in germination medium. (a) Phase-contrast microscopy. (b) Field image using quick-freeze deep-etch replica EM. (c) Quick-freeze and deep-etch replica EM of germinating spore. (d, f) The rodlet layer is marked by a brown asterisk. (e, g) Magnified images of boxed areas in the left panels. Cracks in the rodlet layer (white star) and micro-holes (white triangles) are shown. (h) Field image of quick-freeze and fracture EM. (i, j) Sectioned cell images. Disintegrated core membrane vesicles (white triangles), rodlet layer (brown triangles), and extending core membrane (yellow triangles) are shown.

After being incubated for 4.5 h in the germination buffer (Fig. 6a, b), hair-like structures were still visible in some spores, as shown in Fig. 6c, and the cracks in the rodlet layer became more pronounced (Fig. 6c, d, e, f), with germ cells occasionally emerging from under the cortex. The outgrowth of germinating cells was noted in some spores (Fig. 6g, h). In QFDE-EM, protrusions ranging from 10 to 20 nm in height (n = 11) were observed on the surface of the germinating cell (Fig. 6f, g). At times, filamentous structures enveloping the surface of new vegetative cells were observed (Fig. 6h), indicating that, while the nascent vegetative cells initially showed small protrusions, over time, the cell surface became covered with small filaments.

**Figure 6.**
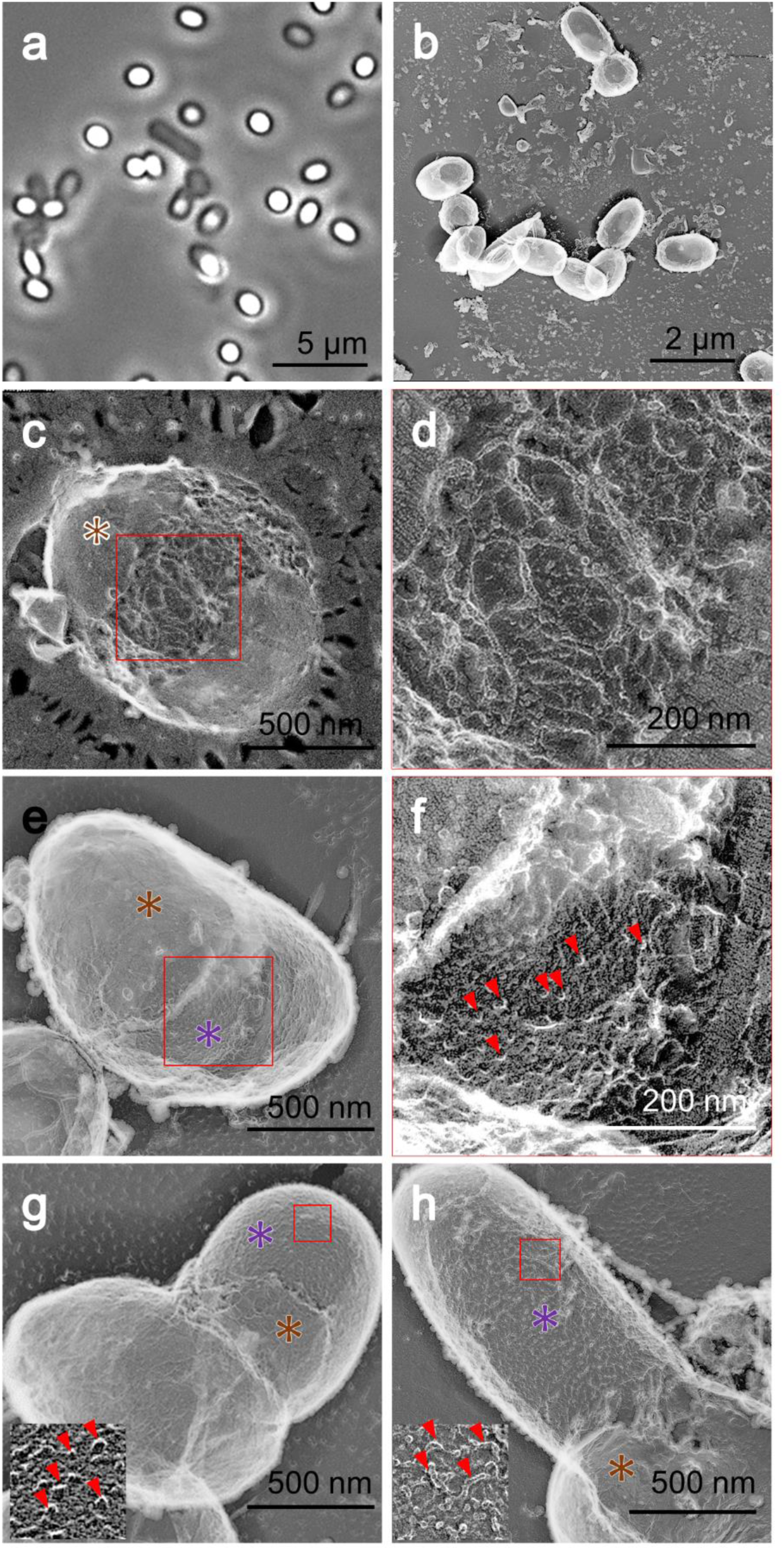
Germinating spores at 4.5 h incubation in germination medium. (a) Phase-contrast microscopy. (b) Field image using quick-freeze deep-etch replica EM. (c, d, e, f) Quick-freeze deep-etch replica EM. (d, f) Magnified images of boxed areas in the left panels. (e) Germ cell surface observed in the crack of the spore. Spore (brown asterisk) and germ cell (purple asterisk) surfaces are shown. (g, h) Partially hatched cells. The remaining spore (brown asterisk) and germinating cell (purple asterisk) are shown. The boxed region is magnified as an inset. Small protrusions are marked by red triangles.

## Discussion

We applied the QFDE-EM technique to *B. subtilis* spores. This method offers several advantages, including the production of high-contrast and three-dimensional images, rapid and chemical-free fixation, and the ability to reveal surface structures with higher resolution, which is not possible using scanning electron microscopy (SEM) [15–17]. The structures we observed were consistent with those reported in previous studies while providing new insights and clearer images.

### Hair-like surface structures

We analyzed the outermost surface of the spores in three dimensions (Fig. 2a–d). Previously identified as a “crust” in thin sectioning studies [8, 9, 26, 27], our observations led us to conclude that the outermost layer consists of thin, hair-like fibers rather than a crust [8]. Attempts to visualize this hair-like surface using the chemically fixed freeze replica technique were unsuccessful [28], likely due to fractures occurring at higher positions without sufficient deep etching. Furthermore, chemical fixation may degrade the quality of the image by potentially crosslinking thin structures to the cell surface [28, 29]

The outermost layer of a spore is composed of proteins encoded by two gene clusters: cotVWXYZ and cgeAB. The hair-like surface structures we observed were flexible, filamentous structures of various lengths and orientations (Fig. 2c, d), suggesting their role in interactions with multiple entities [30–32]. CotV and CotW proteins produced in *Escherichia coli* display helical fibers under EM [33], indicating the involvement of these proteins in the formation of the surface structures [33–36].

### Spike structures

We observed and analyzed spike structures induced by lysozyme treatment (Fig. 2e–h), which had been previously examined using thin sectioning [8]. These structures contained 6-deoxyhexoses, including rhamnose, 3-O-methyl-rhamnose, quinovose, glucosamine, and muramic lactam, a few of which are shared with the PG of the spore [8]. Our images provide a clearer three-dimensional view of these structures (Fig. 2g, h), showcasing their various lengths and orientations. Compared to the hair-like structures, they appear to be more rigid. This suggests their potential role in surface interactions, as previously proposed [8].

### Internal structures

A rodlet layer was identified beneath the hair-like structure (Fig. 3a), consistent with earlier findings reported through chemically fixed freeze replica EM and atomic force microscopy (AFM) [7, 10, 28, 37]. Our observations, which show a periodicity of 8 nm (Fig. 3c), are consistent with these reports. However, we observed that the rodlet layer exhibited two to three overlapping layers, a detail not previously reported (Fig. 3b) [37]. Various genes, including *cotA*, *cotB*, *cotE*, *cotH*, *spoIVA*, and *safA*, are implicated in the formation of the rodlet layer; however, details of its composition remain to be determined [11, 31, 33–35].

Although the outer membrane was identified in a *ponA*-deficient mutant at the same location using cryo-EM [36], its existence in intact mature spores remains to be elucidated [14, 31].

Under the rodlet layer, a porous cortex was identified, as indicated by a pink asterisk in Fig. 3e (view of a section is highlighted by a green asterisk in Fig.3f). The chemical makeup of the cortex PG differs from that of the vegetative cell wall, with 50% of the peptide side chains of N-acetylmuramic acid (NAM) residues being replaced by muramic-δ-lactam [38, 39]. This loosely cross linked layer is presumed to play a role in protoplast dehydration. We then examined the core surface (marked by a red asterisk in Fig. 3e), which appears compact and smooth, aligning with previous AFM observations of *B. anthracis* [13]. Small vesicles were observed in the core membrane (Fig. 3h, i; marked with black triangles), resembling sCMMs (Fig. 3h) [14], which were observed outside the core, indicated by green triangles (Fig. 3g, h, i). This finding is in contrast to previous findings [14]. sCMMs are hypothesized to function as carriers for lipids and proteins, helping the organism withstand environmental stresses such as surfactants, oxygen depletion, starvation, and thermal shock.

### Isolation of peptidoglycan

We successfully isolated the PG sac during vegetative growth by treating it with SDS and protease, as previously demonstrated in *E. coli* [40]. QFDE-EM revealed that the sac displayed a mesh pattern similar to that of the vegetative cell surface, indicating that the PG layer was exposed in vegetative cells (Fig. 4m). These patterns contrast with those of PG isolated from gram-negative bacteria, such as *E. coli*, which exhibited pores [41], likely due to variations in PG layers in terms of thickness and chemical composition.

Although the PG sac was isolated from vegetative cells using the same method, the structure obtained from spores appeared quite opaque under negative staining EM (Fig. 4e). Moreover, the QFDE structure was thicker and rougher than that of the vegetative counterpart (Fig. 4n). This disparity may have stemmed from differences in chemical structures. This hypothesis is further supported by the results of the lysozyme treatment (Fig. 4j, k, l, o), where lysozyme presumably degraded certain susceptible components of the spore PG layer, resulting in the observable porous structure of the spore PG [38, 42, 43].

### Germination

In QFDE images, the hair-like structures remained attached to the spores even after heat treatment, although they were sometimes partially removed due to fracturing (Fig. 5a–c). The initial step in germination appears to be the degradation of the rodlet layer (Fig. 5d– g), which is consistent with previous studies on *B. anthracis* and *Clostridium novyi* NT conducted using AFM [37, 44]. At this stage, sCMMs were not observed, implying their fusion with the core membrane (Fig. 5h–j), as previously described [14]. The thread-like structures in the core membrane may represent the remnants of these vesicles (Fig. 5j).

Under the degrading rodlet layer, we identified a filamentous layer (Fig. 6c, d), which may have been derived from the cortex layer. The germinating cells emerged laterally, in agreement with an earlier study [45]. The small protrusions observed on the core surface (Fig. 6e, f, g) may serve as precursors for the PG filaments observed on vegetative cells (Fig. 6h).

### Concluding remarks

In a previous study, we demonstrated the effectiveness of QFDE-EM in studying spores, including those from the yeast *Saccharomyces pombe* and the bacterium *Actinoplanes missouriensis*, by visualizing multiple layers at different heights with minor modifications. The resistance of spores to washing with pure water is advantageous to preventing unexpected noise [46, 47]. In our study, we successfully applied QFDE-EM to spores of *B. subtilis,* a bacterial spore of significant interest [48]. We hope that our in-depth ultrastructural studies using the QFDE-EM technique will provide valuable insights for future research on bacterial spores.

## Funding

This study was supported by Grants-in-Aid for Scientific Research (A) (JP17H01544) and a JST CREST grant (JPMJCR19S5) awarded to MM.

## Acknowledgments

We thank Junko Shiomi of Osaka Metropolitan University for her technical assistance.

## Data Availability Statements

The data underlying this article are published together.

## Abbreviations

PG: peptidoglycan
CaDPA: Ca^2+^ and dipicolinic acid
EM: electron microscopy
QFDE-EM: quick-freeze deep-etch electron microscopy
LB: Luria-Bertani
PBS: phosphate buffered saline
FFT: fast Fourier transform
QFF-EM: quick-freeze fracture without etching EM
NAM: N-acetylmuramic acid
sCMMs: sub-CM membranes
SEM: scanning electron microscopy
AFM: atomic force microscopy

## Notes

### Competing Interest Statement

The authors have declared no competing interest.

